# Repurposing statins in combination therapy for effective ablation of metastatic breast cancer

**DOI:** 10.1101/2025.05.25.655937

**Authors:** Rifat Aara, Neeha Sinai Borker, Jyothilakshmi Sajimon, Seemadri Subhadarshini, VP Snijesh, Vidya P Nimbalkar, Manju Moorthy, Archana P Thankamony, R Athul Krishnan, Diya Aich, Debolina Das, Mohit Kumar Jolly, Jyothi S Prabhu, Radhika Nair

## Abstract

Metastatic breast cancer (mBC) remains an incurable disease with limited treatment options, highlighting the need for novel therapeutic approaches. Combination therapy with chemotherapeutic agents along with targeted therapy are the most common methods of treatment used in the terminal stages of the disease. Eventual development of resistance to these approaches, leading to fatal outcomes suggests the presence or emergence of a heterogenous population of cells intrinsically resistant to commonly used regimens. Previous work identified a heterogeneous population of metastatic tumor cells with distinct molecular characteristics driven by Macc1 (Metastasis Associated in Colon Cancer 1) overexpression, which could be targeted by lovastatin (transcriptional inhibitor of Macc1). Building on this foundation, the efficacy of lovastatin in targeting metastatic cells with high Macc1 expression was evaluated in lung metastasis, and the regulatory pathways governing Macc1 expression and lovastatin treatment in mBC were investigated. The expression of Macc1, in breast cancer biopsies provides insights into the intra and inter tumor heterogeneity of the Macc1 gene and its correlation with disease outcomes.

While some studies suggest synergistic effects between statins and chemotherapeutic agents, comprehensive evaluations of various combinations and their therapeutic outcomes are still needed. The therapeutic efficacy of lovastatin combined with chemotherapy to determine the most effective treatment regimen that maximizes tumor cell ablation demonstrated that lovastatin can effectively ablate the chemoresistant tumor cells in mBC. The translational implications of this research will identify patient subgroups that may benefit most from statin-chemotherapy combinations that could support the repurposing of statins as cost-effective adjuvant therapies for mBC.

## 1. Introduction

Discovering new therapeutic approaches for metastatic breast cancer (mBC) remains an urgent unmet clinical need. mBC cases are expected to rise dramatically and contribute to most breast cancer-related deaths in the coming years (1). Despite advancements in early detection and treatment, survival outcomes for mBC patients remain dismal, particularly in low- and middle-income countries (2,3). In India, breast cancer patients experience significantly poorer survival rates compared to their counterparts in Western nations. Disturbingly, the disease-free survival rate for stage 4 or mBC is reported to be less than 22%, underscoring the urgent need for more effective treatment strategies and interventions (4).

mBC presents a formidable clinical challenge, primarily due to the lack of effective targeted therapies, rendering systemic chemotherapy to be the cornerstone of disease management. Patients with mBC have already been treated with multiple chemotherapies which have failed, due to the development of drug resistance within the tumor, leading to the advancement of the disease (5). Currently, mBC is managed primarily through combination therapy (5–7). However, using more than one drug in combination is often a case of trial and error in end-stage disease, due to the lack of options left, frail health of the patient as well as the terminal nature of the cancer at this stage.

One of the primary drivers of resistance in mBC is intratumoral heterogeneity (ITH)—the presence of diverse tumor sub-populations with varying degrees of drug resistance (8). This heterogeneity enables selective survival of resistant clones, leading to treatment failure and disease progression (8,9). Therefore, using conventional chemotherapy often proves insufficient, necessitating novel therapeutic strategies to overcome tumor heterogeneity and resistance in mBC. In such cases, identifying and repurposing non-cancer drugs provides a strategic approach by circumventing the time-intensive and expensive process of novel drug discovery and clinical trials. By leveraging their established safety profiles and mode of action, this approach fast-tracks the availability of effective therapies for patients in urgent need (10–13).

In the field of drug repositioning, statins, originally used to treat hyperlipidemia, have demonstrated promising outcomes in breast cancer treatment, supported by epidemiological studies and fundamental research findings (10–16). Clinical cohort studies have demonstrated that patients who use statins, either before diagnosis or following surgery, exhibit a significantly reduced risk of disease recurrence and breast cancer-related mortality compared to non-users (11,12,15–18). Additionally, the anti-cancer potential of statins has been extensively studied through in vitro and in vivo experiments (10,19–21). However, whether statins have clinically meaningful anti-cancer effects in the advanced stage of metastasized tumors remains an area of active investigation, with limited and conflicting findings across various meta-analyses and clinical trials (22,23). Additionally, differences in their impact on primary versus metastatic tumors remain poorly understood, warranting further exploration to optimize their therapeutic potential.

Lovastatin, a lipophilic statin and a competitive inhibitor of 3-hydroxy-3-methylglutaryl-coenzyme A (HMG-CoA) reductase offers several advantages over hydrophilic statins, primarily due to its enhanced ability to penetrate cell membranes, which makes it potentially more effective in improving overall survival and reducing tumor recurrence (24–26). Recent studies identified lovastatin as an effective inhibitor of Macc1 (Metastasis Associated in Colon Cancer 1), a transcriptional regulator of the HGF-MET signaling pathway (27). Previous work in our lab identified Macc1 as a key regulator of metastasis, with Macc1 overexpression linked to poor prognosis and increased resistance among breast cancer patients. We demonstrated that lovastatin, significantly reduced the proliferation rate of aggressive subpopulation of cells with increased metastatic capacity, within the primary tumor. (28). Building on this foundation, we now seek to evaluate the efficacy of lovastatin in targeting metastatic cells with high Macc1 expression in lung metastasis. We demonstrate that lovastatin effectively inhibits cell proliferation and migration in primary tumors as well as lung metastases, with a more pronounced reduction in Macc1 expression observed in the lung metastases. We next elucidate the molecular mechanism of lovastatin in mBC by identifying and validating the signaling pathways involved in metastatic cells.

The combinatorial effect of lovastatin with chemotherapeutic agents in breast cancer remains controversial, with studies reporting conflicting results. Some research suggests that lovastatin exhibits synergistic effects, enhancing the sensitivity of chemotherapy and overcoming resistance (29–31). In contrast, few studies have reported no significant or even antagonistic effects when lovastatin is combined with certain chemotherapeutic agents (32,33). Given these contradictory findings, further comprehensive investigation is crucial to delineate the conditions under which lovastatin can be effectively integrated into mBC treatment, thereby establishing an optimized therapeutic strategy. To address therapeutic discrepancies and establish the efficacy of combinatorial treatment in a metastatic setting, we have evaluated the effect of lovastatin in combination with standard chemotherapy used for treating mBC patients. The results of this work will set the stage for the rationale design of lovastatin and chemotherapy as a strategy to target mBC.

## 2. Results

### 2.1 Metastatic tumors marked by high Macc1 expression and heterogenous expression patterns

In our previous work, we identified a distinct metastatic population of cells from the primary mammary tumor (T1), driven by high Macc1 expression (28). We demonstrated that lovastatin, a cholesterol-lowering agent and a transcriptional inhibitor of Macc1, effectively targets these highly metastatic cells. In our current work, we used spatial profiling to investigate the distribution and heterogeneity of MACC1 expression in clinical samples of breast cancer patients. We first examined the mRNA level of Macc1 across the 50 breast tumors in all the regions (ROI) selected using the data from GeoMx WTA panel. **Figure 1A** shows the normalized expression of MACC1 across selected representative tumor cores. Expression levels of MACC1 varied across the tumor tissues, which showed differential expression within distinct tumor areas indicating both inter and intra-tumoral heterogeneity of MACC1 among the breast cancer tissue. The association of MACC1 expression across different clinicopathological features of the breast tumors showed MACC1 expression was significantly higher in Grade 3 tumors (**Figure 1B**) suggesting its correlation with aggressive tumor features. Moreover, MACC1 expression was elevated in triple-negative breast tumors (TNBC N=20) compared to hormone receptor-positive (HR+ N=25) tumors (**Figure 1A**, **C**). Given this strong association between MACC1 and aggressive tumor features, we next investigated its role in the metastatic setting. We isolated phenotypically heterogeneous tumor cells from matched lung metastases of the 4T1 GFP (Bulk) murine model and focused on the L1 subpopulation, characterized by adherent morphology. L1 cells exhibited a significantly higher proliferation rate compared to both Bulk and T1 cells, along with an increased proportion of cells in the S phase of the cell cycle **(Figure S1A, B**). Additionally, L1 cells also displayed elevated Macc1 expression compared to the T1 and Bulk tumor cells (**Figure 1D**).

**Figure 1.**
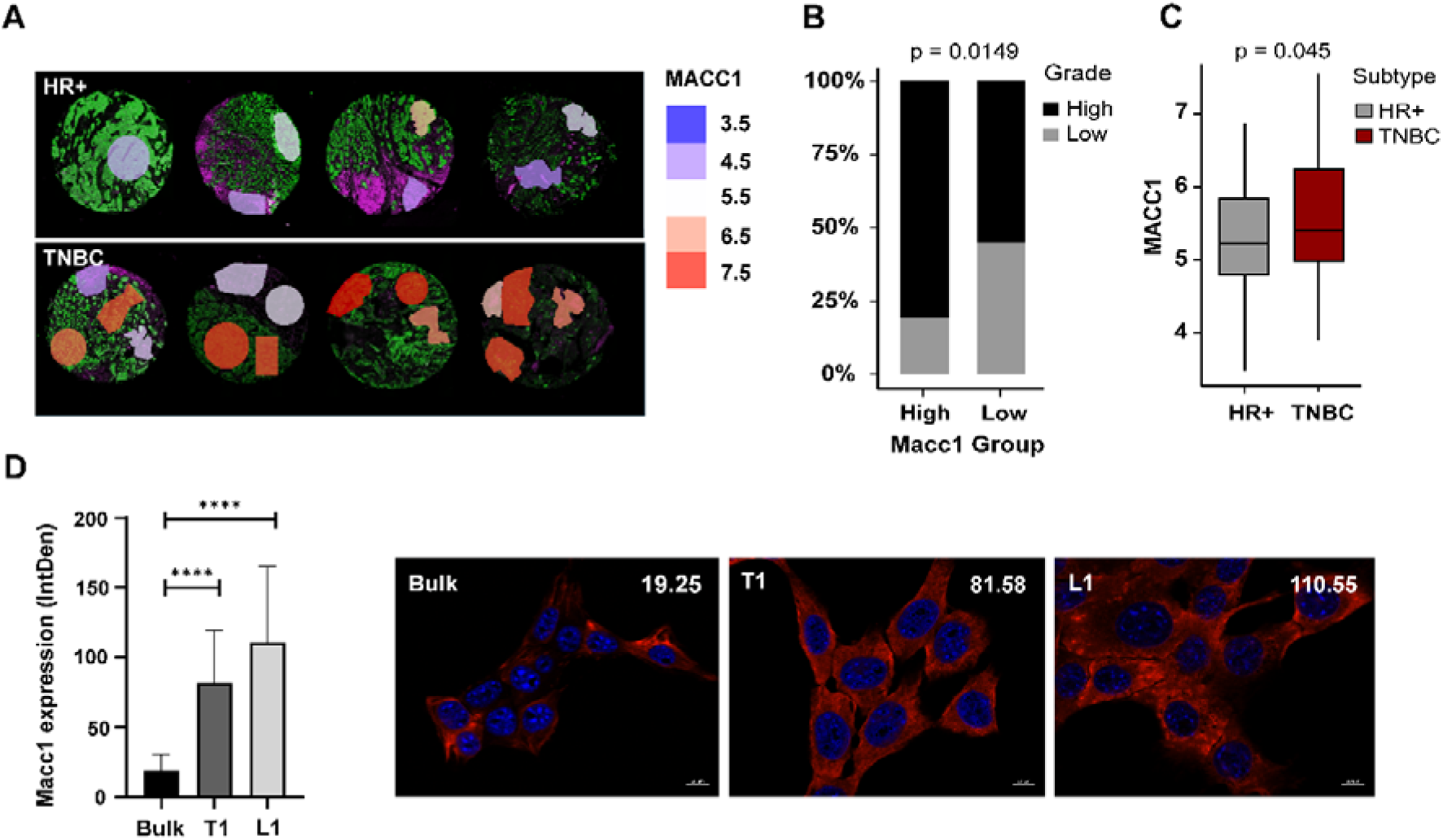
Spatial and quantitative analysis of MACC1 expression in metastatic tumor. A) Spatial profiling of mRNA expression of MACC1 across selected representative tissue microarray cores of hormone receptor-positive (HR+) and triple-negative breast cancer (TNBC) tumors done by GeoMx platform showing the inter and intra-tumoral heterogeneity in its expression. Log2 Q3 normalized values represented here for MACC1 using SpatialOmicsOverlay R package. B) Distribution of histological tumor grade across MACC1 high and low tumors (using the mean values as cut-off) showing a higher proportion of high-grade tumors in tumors with increased expression of MACC1. C) Expression of MACC1 (log2 normalized values) across the Hormone receptor-positive (HR+) and triple-negative breast cancer tumors (TNBC), showing higher expression in TNBC. D) Quantification of Macc1 expression in Bulk, T1, and L1 cells using Immunofluorescence assay. L1 cells have significantly higher MACC1 expression compared to the Bulk and T1 cells. Representative images of Macc1 expression in Bulk, T1, and L1 cells. Red and blue represent Macc1 and DAPI staining respectively. Images were taken using Zeiss Apotome fluorescence microscope at 63x magnification with a scale bar corresponding to 10 µm.

### 2.2 Targeting heterogeneous metastatic tumor cells using Lovastatin

We have previously shown that treatment with lovastatin, which is a Macc1 transcriptional regulator, led to the ablation of T1 cells via downregulation of Macc1. We hypothesized that lovastatin would also be effective on the L1 lung metastatic cells, which have higher Macc1 expression than the T1 cells. In order to test this hypothesis, we assessed the cytotoxic effect of lovastatin on Bulk, T1, and L1 tumor cells. We observed a dose-dependent reduction in cell viability, with significantly lower IC50 values of 4.9 μM and 4.6 μM for T1 and L1 cells respectively, compared to 8.7 μM in the Bulk tumor cells (**Figure 2A**).

We decided to use the IC50 concentration of lovastatin of the Bulk cell type (8.7 μM) for further experiments, as it would replicate the concentration of a drug that is given based on the bulk tumor in the clinic. We assessed the viability and Macc1 expression in Bulk, T1, and L1 cells after 48 hours of lovastatin treatment. There was a significant reduction in cell viability in the T1 and L1 cells when compared to the heterogenous Bulk cells. This result indicates that the T1 and L1 cells are more sensitive to lovastatin treatment **(Figure S1C, D, E)**.

We next assessed whether lovastatin modulates Macc1 expression in lung metastases. Macc1 levels in untreated (-) and lovastatin-treated (+) groups of Bulk (B^-/+^), T1 (T1^-/+^), and L1 (L1^-/+^) cells were analyzed at single-cell level using immunofluorescence followed by quantification through integrated density analysis. Macc1 expression was significantly downregulated in lovastatin-treated L1 cells compared to the untreated group, with a more pronounced reduction than in Bulk and T1 cells (**Figure 2B, C, Figure S2A)**. Scatter plot analysis of untreated and lovastatin-treated groups from Bulk, T1, and L1 cells revealed a significant shift in MACC1 expression. We binned the cells based on the range of Macc1 expression and found more than 95 % of the treated cells in the T1 and L1 population shifting to the lowest Macc1 expression interval (0–50) **(Table S1, Figure S2 B, C, D)**. This clearly shows the effect of lovastatin on Macc1 expression and subsequent downstream effects observed on the metastatic phenotypes driven by Macc1.

Based on previous studies which have suggested a dose-dependent effect of lovastatin in the cell cycle, we next looked at the effect of lovastatin on the cell cycle in the heterogeneous metastatic tumor cells. We analyzed the cell cycle phases in Bulk, T1, and L1 cells with lovastatin treatment (**Table 1**). Interestingly, we observed a change in only the T1 group upon lovastatin treatment, with a significant increase in the sub G0 phase **(Figure S3A, B, C)**. We further found that cell cycle-related genes like Cdkn1a, p53 and Ccne1 were significantly altered only in the Bulk cell population **(Table S2, Figure S3D)**.

Additionally, lovastatin treatment markedly inhibited the migratory ability in Bulk, T1, and L1 cells **(Figure S4A, B, C)**. We also observed a reduction in the area of the nucleus of T1 and L1 cells compared to the untreated group **(Figure S4D)**, which we are investigating further.

**Figure 2.**
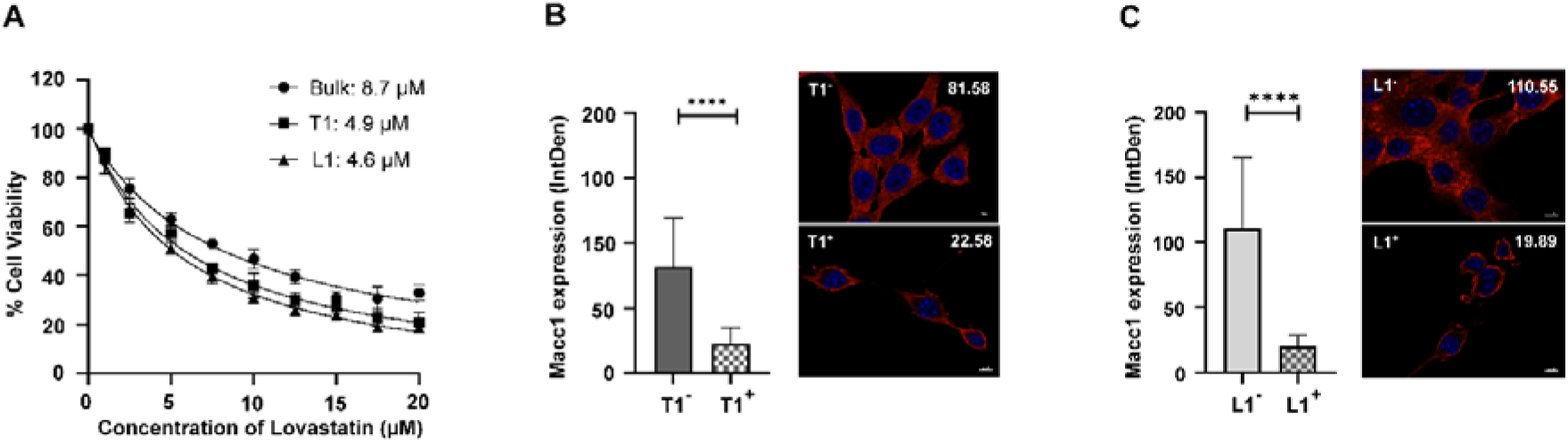
Effect of lovastatin on metastatic tumor cells. A) Graph showing dose-dependent effect of lovastatin on the cell viability of Bulk, T1 and L1 cells treated for 48 hours measured using MTT assay. IC50 values of lovastatin in Bulk, T1 and L1 cells were calculated using GraphPad Prism software (n=3). Effect of Lovastatin (8.7 µM concentration) on the Macc1 expression of B) T1 cells and C) L1 cells upon 48 hours of treatment at quantified using Immunofluorescence assay. T1 and L1 cells treated with Lovastatin showed a significant reduction in Macc1 expression compared to untreated cells. Immunofluorescence images of Macc1 expression in the untreated vs Lovastatin-treated T1 and L1 cells.

**Table 1.**
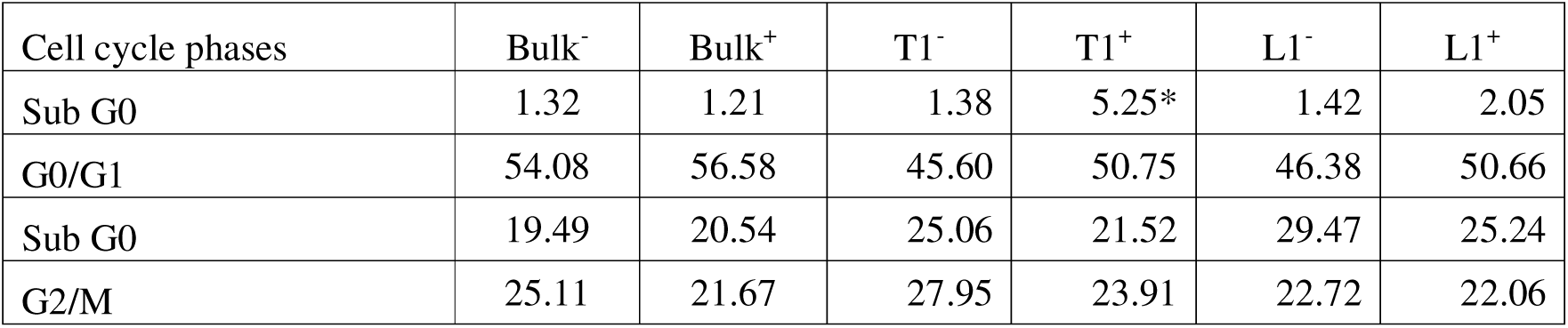
Effect of lovastatin on the cell cycle. Results are expressed as the percentage of Bulk, T1 and L1 cells in different cell cycle phases. Regulation of the phases is with respect to treated vs control. All experiments were performed in three biological replicates and data were expressed as mean ± standard deviation. Two-way ANOVA statistical test was used. *p < 0.05, **p < 0.01, ***p < 0.0001.

### 2.3 Molecular basis of lovastatin treatment on metastatic cells

We next looked at the molecular players involved in the effect of lovastatin on the metastatic breast cancer cells. We looked at the transcript levels of Macc1, Met, Wnt7a, Pikr3r and Map, as well as EMT (Cdh1, Vim, Tgfβ2) and Dormancy (Nr2f1) markers to discern the epithelial and quiescence state of the heterogenous tumor cells **(Table S3)**.

There was a significant decrease in Macc1 expression in all cell groups upon lovastatin treatment (**Figure 3A**). Analysis of the epithelial and dormancy markers revealed a significant upregulation in the mesenchymal marker Vimentin in the Bulk and T1 cells (**Figure 3B**), and an upregulation in the epithelial marker, Cdh1 only in Bulk cells suggesting that epithelial transition mechanisms may be primarily restricted to the bulk population and are not prominently active in sub-populations, T1 and L1 under treatment conditions **(Figure S5A)**. There was a downregulation in Tgfβ2 in the lovastatin treated samples of all three cell groups, which was most pronounced in L1 (**Figure 3C**), which could correlate with changes seen in EMT and migratory phenotypes in the L1 cells. We noted a significant decrease in the dormancy gene Nr2f1 in L1 treated cells, in contrast to the increase in the Bulk cell population upon lovastatin treatment (**Figure 3D**). Interestingly, other genes like Map and Pik3r3, with a potential role in Macc1 regulation did not significantly change in the metastatic populations suggesting that this signaling axis may not be a major contributor to Lovastatin response in these sub-populations **(Figure S5B, C)**. This indicates a potential new regulatory role of Macc1 in metastatic sub-populations, T1 and L1 cells which we are currently exploring.

We also looked at transcription factors like Sp1 which play a role in the transcriptional regulation of Macc1(34,35). Molecular docking was conducted to assess the potential interaction between Sp1 and lovastatin, focusing on its zinc finger domains (**Figure 3E, F, G)**. C2H2-type zinc fingers coordinate Zn²□ ions via conserved histidine and cysteine residues, which are essential for maintaining structure and enabling DNA binding. The high-confidence AlphaFold model of Sp1’s zinc finger region was docked with lovastatin. The docked complex with the lowest Vina score (–7.2 kcal/mol) revealed interactions with key residues: LYS635, TYR637, HIS642, ALA645, HIS646, TRP649, ARG654, ARG666, PHE667, THR668, ARG669, ASP671, GLU672, and ARG675. The zinc finger domain is critical for Zn²□ coordination and DNA binding, suggesting that lovastatin may interfere with Sp1’s structural integrity and function.

**Figure 3.**
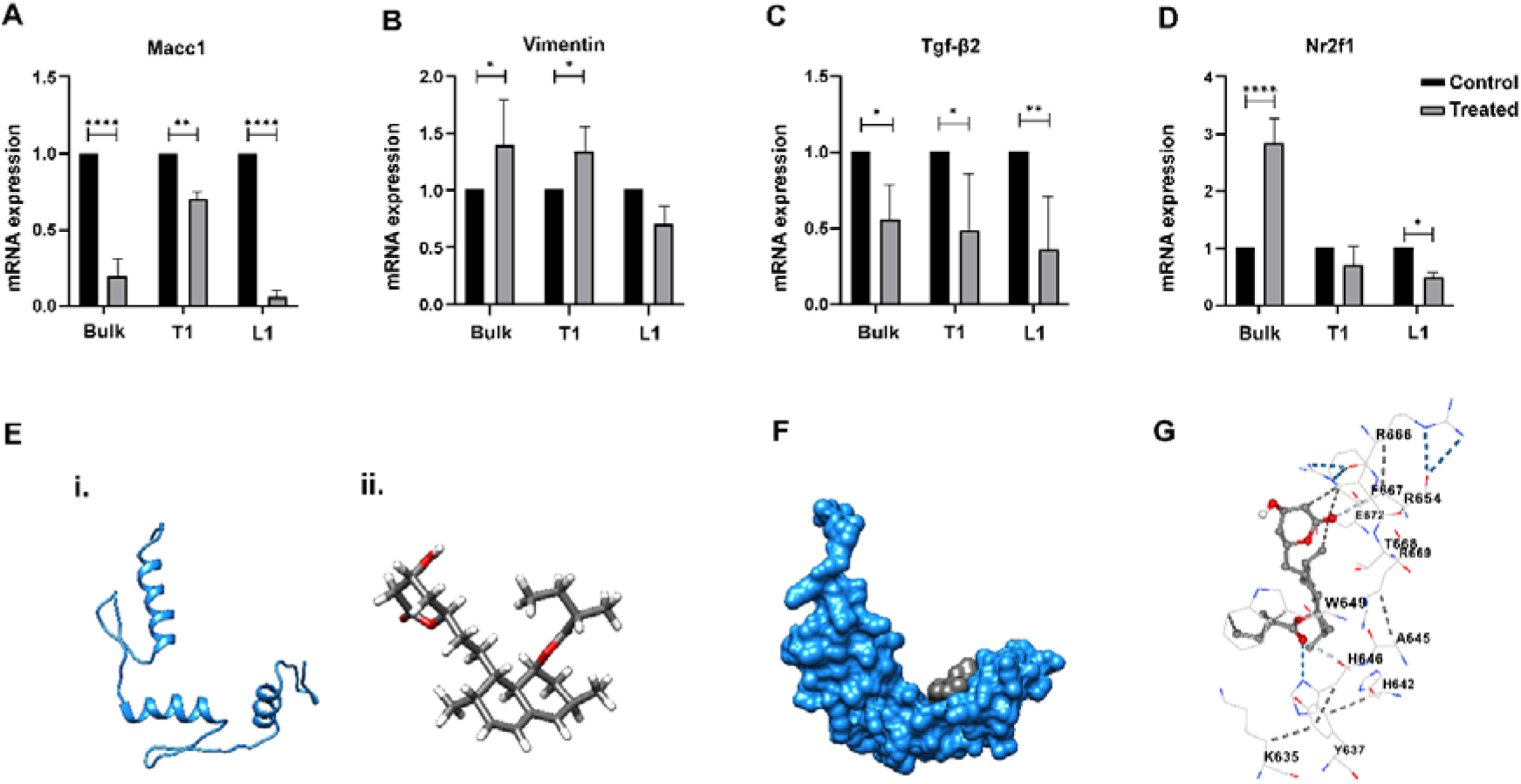
Molecular changes with lovastatin treatment on metastatic cells. The mRNA levels of A) Macc1 B) Vimentin C) Tgf-β2 and D) Nr2f1 was analyzed by qRT-PCR, post 48 hours of lovastatin treatment, normalized with β-Actin. Macc1 expression is significantly downregulated upon treatment in the Bulk population, T1 and L1 cells. Vimentin, a mesenchymal marker, shows a significant reduction in expression in the Bulk and T1 cells post-treatment, with no significant change in the L1 cells. Nr2f1, a marker associated with dormancy, is significantly upregulated in the bulk population upon treatment, while expression is moderately increased in the L1 cells. The data was normalized to β-actin and analyzed using 2^-ΔΔ*CT*^ method. All the experiments were performed in three biological replicates and the data were expressed as mean ± standard deviation. Two-way ANOVA statistical test was used. *p < 0.05, **p < 0.01, ***p < 0.0001. E) Structural Overview: (i) Cartoon representation of the AlphaFold-predicted structure of Sp1 (AF-P08047-F1-v4), filtered to include high-confidence residues (pLDDT = 70) spanning positions 625–719, corresponding to the three C2H2 zinc finger motifs. (ii) Stick representation of the ligand, lovastatin, highlighting its spatial orientation prior to docking. F) Surface view of the docked Sp1-lovastatin complex, illustrating the spatial fit of the ligand within the predicted binding pocket. G) Three-dimensional visualization of the protein-ligand interactions within the receptor cavity. Dark blue lines denote strong hydrogen bonds, light blue lines indicate weak hydrogen bonds, and grey lines represent hydrophobic interactions.

### 2.4 Synergistic effect of combination therapy of lovastatin and chemotherapy on metastatic tumor cells

Finally, as proof of concept, we tested if we could target chemoresistant metastatic cells by concurrent combination therapy with lovastatin. We first determined the IC50 of the commonly used chemotherapies, 5-fluorouracil (5-FU) and Paclitaxel in Bulk, T1 and L1 cells (**Table 2**). We observed a significant increase in the IC50 for 5-FU in the T1 and L1 cells compared to the Bulk population, indicating that T1 and L1 cells were more resistant to 5-FU (**Figure 4 A)**. The T1 and L1 cells were more sensitive to Paclitaxel **(Figure S5D)**. We next combined Lovastatin (Bulk IC50-8.7 μM) with 5-FU (Bulk IC50-0.26 μM) in concurrent combination treatment for 48 hours and checked the viability of each cell type with monotherapy (L represents Lovastatin monotherapy and F represents 5-FU monotherapy) and combination therapy (L+F). Strikingly, we saw a synergistic effect on the viability of lovastatin and 5-FU treatment, which was significant in the metastatic T1 and L1 cells, when compared to monotherapy (L, F) and untreated group (C), (**Figure 4B, C, Figure S5E)**. This strongly suggests that lovastatin can sensitize metastatic cells (T1 and L1), which are otherwise resistant to 5-FU, thereby potentially preventing relapse associated with 5-FU monotherapy (**Figure 5**).

**Figure 4.**
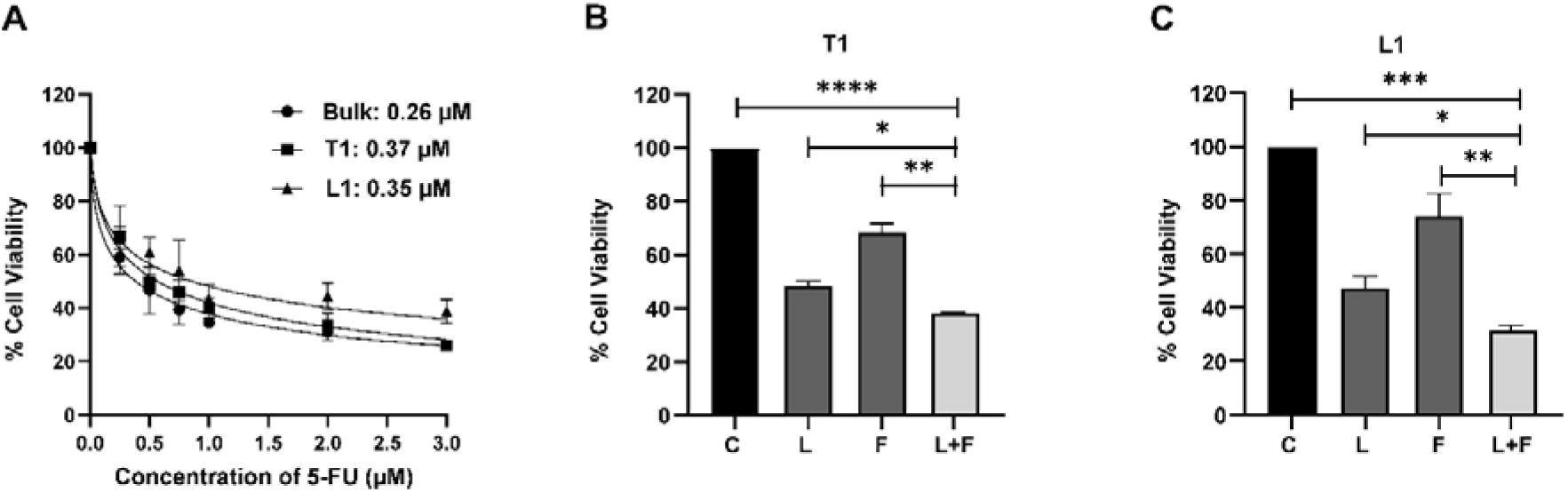
Targeting Bulk, T1 and L1 tumor cells using combination therapy. A) Graph showing dose-dependent effect of 5-FU on the cell viability of bulk, T1 and L1 cells treated for 48 hours measured using MTT assay. IC50 values of 5-FU in Bulk, T1 and L1 cells were calculated using GraphPad Prism software. T1 and L1 cells showed higher resistance to 5-FU treatment compared to the Bulk. Graphs representing the effect of 5-FU in combination with lovastatin (at the concentrations corresponding to the IC50 of 5-FU and lovastatin in the Bulk tumor cells) on B) T1 and C) L1 cells. There is a significant reduction in cell viability in the cells treated with 5-FU in combination with lovastatin compared to cells treated with only 5-FU and lovastatin and untreated group in T1 and L1 cells. Statistical significance was calculated using Unpaired t test with Welch’s correction with * p value < 0.05 and ** p value <0.001.

**Table 2.** IC50 values of Lovastatin, 5-FU and Paclitaxel on Bulk, T1 and L1 cells.

| Cell type/Drug | Lovastatin ($\mu\text{M}$) | 5-FU ($\mu\text{M}$) | Paclitaxel (nM) |
| --- | --- | --- | --- |
| Bulk | 8.7 | 0.26 | 69.3 |
| T1 | 4.9 | 0.37 | 48.7 |
| L1 | 4.6 | 0.35 | 36.2 |

**Figure 5.**
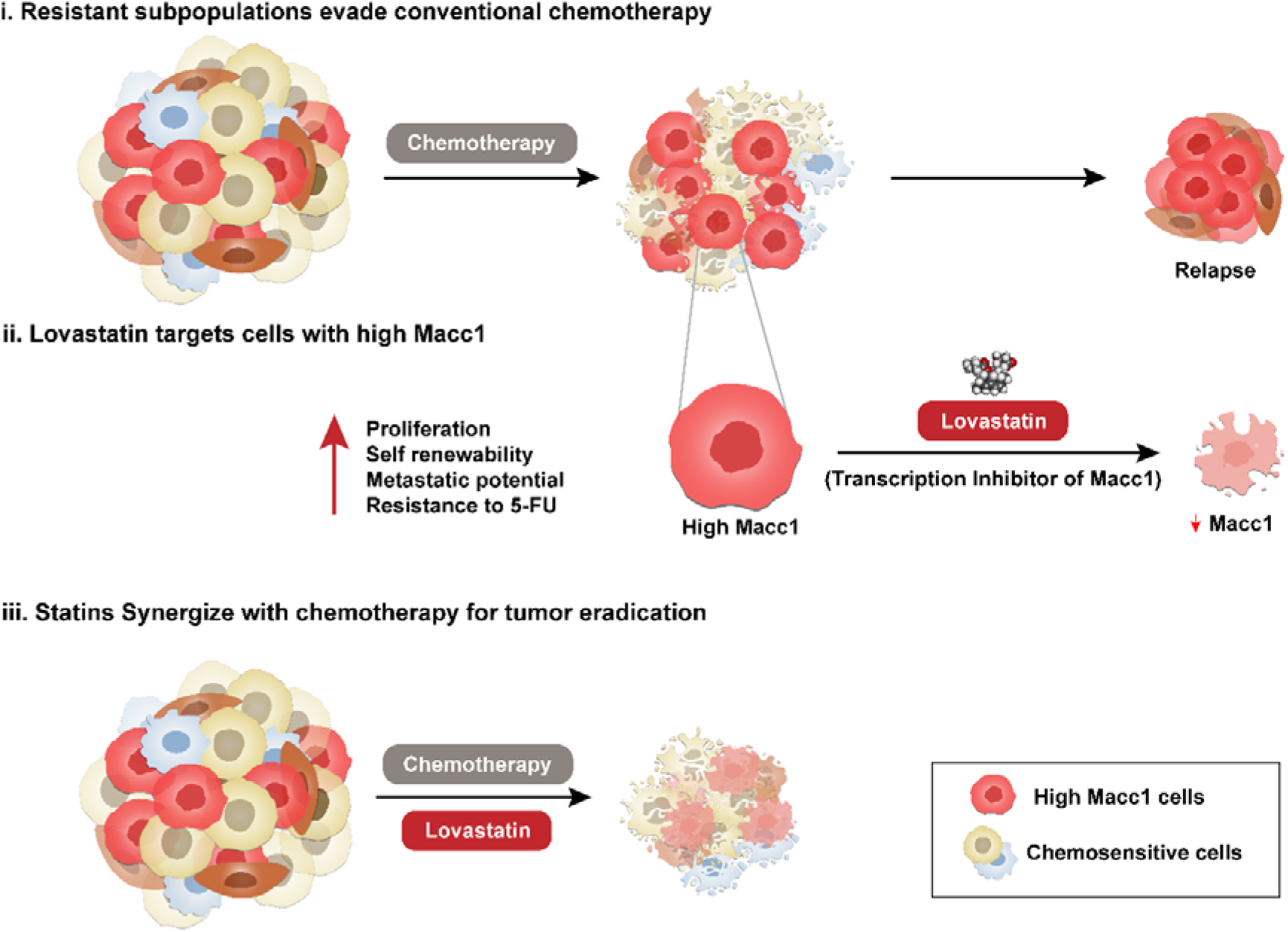
Enhancing chemotherapeutic efficacy through statin-driven tumor ablation. Metastatic tumors consist of heterogeneous cell populations, including a distinct subpopulation of cells with high expression of Macc1. Conventional chemotherapy primarily targets the chemosensitive tumor cells (blue), but fails to eliminate resistant subpopulations, including those with high Macc1 expression (red). These surviving cells persist and drive recurrence and metastasis. Lovastatin, which is a clinically approved statin, functions as a transcriptional inhibitor of Macc1, leading to downregulation of its expression which reduces the survival advantage of Macc1-overexpressing cells. The combined treatment of lovastatin with chemotherapy effectively overcomes tumor heterogeneity by targeting both chemosensitive and chemoresistant Macc1-high cells, leading to more effective tumor ablation.

## 3. Discussion

This study provides compelling evidence for the potential of lovastatin as a targeted therapeutic agent in metastatic breast cancer (mBC), particularly in the context of tumor subpopulations from lung metastases characterized by high Macc1 expression. Our findings build upon earlier observations that Macc1, a key regulator of metastasis and tumor aggressiveness, is markedly elevated in lung metastatic cells and can be effectively suppressed by lovastatin (28). We demonstrate that lovastatin not only inhibits the proliferation of these aggressive subpopulations but also significantly downregulates Macc1 expression at both transcript and protein levels, with a more pronounced effect in metastatic (L1) cells compared to primary tumor-derived populations.

Numerous studies in the past have evaluated the expression of both RNA and protein levels of MACC1 in primary treatment naïve invasive breast tumors and have shown the varied expression amongst subtypes of breast cancer (36–39). Ali et al examined the mRNA expression of MACC1 in 120 tumors of invasive breast tumors and shown its correlation with aggressive tumor characteristics, differential tumor microenvironment and poor prognosis (36). Our results are not only in agreement with these studies in showing intertumoral heterogeneity of MACC1 expression, spatial profiling of mRNA also showed significant differences in intra tumoral expression of MACC1. While direct spatial profiling studies of MACC1 in breast tumors are limited, the observed variability in our results though in limited number of tumors, across different tumor subtypes and clinical features suggests potential spatial heterogeneity in its distribution.

In line with previous studies linking MACC1 to poor prognosis and resistance in breast cancer (36–38), our data suggests that the therapeutic efficacy of lovastatin may be partly mediated by its ability to target MACC1-driven molecular programs. The observed altered cell morphology, reduction in migratory capacity and nuclear area in the T1 and L1 cells further highlights the pleiotropic effects of lovastatin and opens new avenues to explore its broader impact on metastatic cells.

We investigated whether lovastatin treatment could affect the cell cycle regulation in these cell lines as previous studies have shown a dose-dependent effect of lovastatin, particularly a G1 phase arrest in breast cancer cell lines (40–43) . While we observed a significant shift in the sub G0 phase in the T1 cells, Bulk and L1 cells did not show any significant changes in any cell cycle phases upon Lovastatin treatment. Our study showed no evident cell cycle arrest, indicating the lovastatin may not act through cell cycle pathway.

At the molecular level, our study sheds light on potential pathways through which lovastatin exerts its anti-metastatic effects in breast cancer. A central observation was the consistent and significant downregulation of Macc1 across all cell populations following treatment, underscoring its role as a target of lovastatin. Given that Macc1 is a known transcriptional regulator of the HGF-MET signaling axis, which promotes proliferation, migration, and metastatic dissemination, its suppression likely disrupts key survival and motility programs in metastatic cells (44–46) . Differential modulation of genes associated with epithelial (Cdh1), mesenchymal (Vimentin), and dormancy (Nr2f1) programs revealed cell type specific effects. Notably, the reduction in Nr2f1 in T1 and L1 cells, contrary to its increase in the bulk population, could reflect a loss of dormancy-related survival advantages in these aggressive subpopulations under statin pressure, warranting further investigation into how dormancy pathways intersect with statin responsiveness.

Beyond MACC1, our transcriptomic analysis revealed downregulation of Tgfβ2 in all three cell populations, with the most pronounced effect in L1 cells. Tgfβ2 is a cytokine implicated in tumor progression, immune evasion, and epithelial-mesenchymal transition (EMT). Its inhibition suggests that Lovastatin may impact both cell-intrinsic signaling and potentially contribute to impaired cell cycle progression, as Tgfβ signaling is known to regulate G1/S transition and maintain a pro-survival transcriptional program(47).

We also observed cell type specific regulation of other key pathways. For instance, Wnt7a, Pik3r3, and Mapk-associated transcripts were differentially modulated, particularly in the bulk population, suggesting that lovastatin’s mechanism of action may vary across heterogeneous tumor subpopulations. The discrepancies observed across subpopulations in terms of pathway modulation and phenotypic outcomes highlight the underlying complexity of metastatic disease. While Macc1 appears to be a common regulatory node, the context-dependent responses such as selective changes in cell cycle progression, EMT, and dormancy signaling underscore the presence of distinct molecular circuits within metastatic clones. These differences also suggest that lovastatin may engage alternate or compensatory mechanisms depending on the cellular context.

The transcriptional regulation of MACC1 plays a pivotal role in promoting tumor aggressiveness and metastatic potential, making it a valuable therapeutic target across various cancers. Previous works have identified the core promoter region of MACC1 and demonstrated that transcription factors such as Sp1, AP-1 (c-Jun), and C/EBP directly bind to this promoter to regulate its expression (27,48,49). Disrupting the interaction of these transcription factors significantly impaired MACC1 expression Further extending these findings, a high-throughput screen conducted by the same group showed that small-molecule inhibitors like Rottlerin and lovastatin are capable of disrupting MACC1 transcription by inhibiting the binding of these transcription factors to the MACC1 promoter, thereby downregulating MACC1 expression.

Building on this foundational understanding, our study reinforces the critical role of Sp1 in MACC1 regulation focusing on Lovastatin. Molecular docking analyses revealed that lovastatin interacts with key residues in the zinc finger domain of Sp1, potentially disrupting its DNA-binding ability and transcriptional activity. Given the established role of Sp1 as a major activator of MACC1 transcription, such interference could plausibly contribute to the observed reduction in MACC1 expression following Lovastatin treatment. However, further experiments are required to validate if this interference translated into a significant downregulation of MACC1 in metastatic tumor subpopulations.

Given that most mechanistic studies on statins in cancer have focused on primary tumors (50–52), there is limited understanding of their role in metastasis-specific biology. Our findings emphasize the need for deeper and more extensive analyses, particularly in breast cancer models focused on metastasis. Elucidating metastasis-specific signaling rewiring and identifying consistent molecular vulnerabilities will be essential for optimizing statin-based strategies and overcoming treatment resistance in the clinical setting.

mBC remains a formidable clinical challenge due to its poor prognosis and limited treatment options. Unlike early-stage disease where surgery, radiation, and targeted therapies offer curative potential, metastatic disease is largely managed through systemic chemotherapy, which remains the primary therapeutic modality. However, its effectiveness is often short-lived due to the emergence of intrinsically resistant tumor subpopulations, driven by underlying intratumoral heterogeneity. In this context, combination therapy has become a cornerstone strategy to enhance treatment efficacy. Yet, in the clinical setting, combinations are often chosen empirically, guided by trial-and-error rather than mechanistic understanding. This underscores the pressing need for rational, biology-driven combinations based on the molecular characteristics of resistant subpopulations, to improve outcomes and minimize toxicity.

Importantly, our findings offer new insights into the potential of combination therapy of lovastatin and chemotherapeutic agents to overcome drug resistance in mBC. To evaluate the ability of Lovastatin to enhance chemotherapeutic efficacy, we first assessed the sensitivity of Bulk, T1, and L1 cells to two commonly used agents in the clinical management of mBC: Paclitaxel, and 5-FU. Interestingly, T1 and L1 cells exhibited more resistance to 5-FU, as reflected by higher IC50 values, when compared to the Bulk population.

This differential response provided a compelling rationale to specifically investigate 5-FU in combination with lovastatin, to test if combination therapy would be more effective. Concurrent treatment with lovastatin and 5-FU led to a synergistic effect and significant reduction in viability of T1 and L1 cells, than with monotherapy. Importantly, this effect was achieved using drug concentrations corresponding to the IC50 values for the Bulk population, highlighting that lovastatin sensitizes intrinsically resistant subpopulations without requiring increased drug dosages.

Further in vivo studies and expanded mechanistic analyses will be essential to validate these findings and guide rational design of personalized statin-based combination therapies in metastatic breast cancer. From a therapeutic standpoint, this result holds promise for strategically integrating statins with existing chemotherapeutics to selectively target resistant clones while minimizing systemic toxicity. Given its favorable safety profile, repurposing Lovastatin in mBC offers a low-cost, readily translatable option.

## 4. Conclusion

Our study provides compelling evidence that lovastatin, a widely used cholesterol-lowering drug, has great potential to be repurposed as a treatment regimen for mBC. By targeting MACC1, a key driver of metastasis and chemoresistance, we have shown that lovastatin inhibits cell proliferation and downregulates MACC1 expression at transcript and protein levels, with a more pronounced effect in metastatic cells. We also showed that combining lovastatin with chemotherapy like 5-FU effectively overcomes tumor heterogeneity by targeting both chemosensitive and chemoresistant Macc1-high cells, leading to more effective tumor ablation. Our findings offer a proof of concept of repurposing lovastatin as an affordable and time-efficient option when used alongside current chemotherapy treatments. This approach can help meet the urgent need for better treatment strategies in mBC. Overall, our work lays the foundation for future clinical trials using lovastatin-chemotherapy combinations which will help develop more effective treatments for people living with this aggressive and currently incurable cancer.

## 5. Methods

### Cell culture

Bulk, T1 and L1 cells were cultured in DMEM/F12 (Invitrogen, 11330-032), supplemented with 10% FBS (Thermo Fisher Scientific, 10270-106) and 1% Penicillin (100 units mL^-1^) /Streptomycin (100 µg/mL) (Thermo Fisher Scientific, 15070-063). The cells were grown at 37^°^C in a 5% CO_2_ incubator.

### Cell proliferation assay

Cells were seeded into 96 well plates in triplicates and cell proliferation were measured for 24, 48 and 72 hours using MTT (Thermo Fischer Scientific, M6494) assay. 10μl MTT solution (5mg/ml) was added to each well and incubated for 2 hours at 37°C. After incubation, the media was removed, and the purple-colored formazan crystals were dissolved by adding 100 μl DMSO was added to each well. The absorbance was measured at 570 nm using Emax ® Plus Microplate Reader.

### Spatial profiling

Spatial profiling on the breast tumors was performed by Digital Spatial technology (DSP) using the Nanostring Geo MX Whole Transcriptomic Panel (WTA). These tissues were chosen from breast cancer cohort collected after ethical approval from institutional ethical committee. Tissue micro array (TMA) was constructed using selected tumor areas (1.5mm core size) from 50 formalin fixed paraffin embedded (FFPE) blocks. Briefly, 5μm FFPE tissue sections were mounted on Superfrost® Plus Micro Slides (Invitrogen, 1255015) and incubated at 65°C for an hour. Sections were deparaffinized, rehydrated, and subjected to antigen retrieval in pH 9.0 solution (Invitrogen, 2450723) at 100°C for 15 minutes, followed by proteinase K treatment. Hybridization involved overnight incubation with WTA panel (NanoString, 18000 genes) at 37°C. Post-hybridization washes included SSC buffer and formamide washes. Tissues were then blocked in Buffer W and stained with morphological markers, including PanCK and SYTO 13, for identifying epithelial cells and DNA, respectively. Regions of interest (ROIs) were selected based on tissue morphology, automatic segmentation using the PanCK marker distinguished tumor (PanCK+) from non-epithelial regions (PanCK-). Indexing oligonucleotides from each Area of illumination (AOI) were released under UV light and collected. For library preparation and sequencing, the oligonucleotides were dried, reconstituted, and sequenced using the Illumina NovaSeq 6000 platform, with 2 x 151 base paired end reads. The entire process ensured high-quality data collection for subsequent analysis. The sequenced fastq files were processed further to convert into dcc (Digital Count Conversion) format using the GeoMx NGS Pipeline v.2.3.3.10 from NanoString for each Region of interest (ROI) and analysed using GeoMxTools R package (v3.8.0). QC for both probe and segment, normalization and unsupervised clustering was performed using GeoMxTools. After appropriate QC on segments and target genes, the counts were normalized using the Q3 method of normalization. Spatial overlaying of gene expression over the tumor cores were performed using SpatialOmicsOverlay (v1.8.0) R package.

### Drug treatment

Lovastatin (Santacruz, sc-200850), 5-Fluorouracil (Sigma-Aldrich, F6627) and Paclitaxel (Sigma-Aldrich, T7191) was dissolved in dimethyl sulfoxide (DMSO). Stocks were stored at -20°C in aliquots to avoid repeated freeze thawing. To see the effect of drugs on cell proliferation, cells were seeded in a 96 well plate. After 24 hours, the cells were treated with increasing concentrations of the drug, with a corresponding concentration of DMSO less than 0.1%. The cell proliferation 48 hours post lovastatin treatment was analyzed by MTT assay. IC50 was calculated using GraphPad prism software.

### Migration assay

Cells were seeded in 24-well plates and cultured in medium containing 10% FBS to near confluence of the cell monolayer. The cells were carefully scratched using a sterile 10 µL pipette tip, and cellular debris was removed by washing with media. The wounded monolayer was incubated with or without Lovastatin for 24 hours. Cell migration into the wound area was photographed using microscope.

### Immunofluorescence assay

Cells were seeded in a 24-well plate (Thermo Fisher Scientific, 142475) containing sterilized coverslips (Blue star, microscopic cover glass circular 12 mm, 10 gm). The cells were washed with PBS and fixed in 4% PFA (Himedia, TCL119) for 10 minutes. After fixation the cells were blocked and permeabilized using 2% FBS (Thermo Fisher Scientific, 10270-106) in 0.3% PBS-Tween (Sigma-Aldrich, P1379) for 30 minutes. The cells were thoroughly washed and stained with primary antibody (Macc1, Santa Cruz Biotechnology, 5179) overnight at 4°C followed by incubation with Donkey anti-Rabbit IgG (H+L) Highly Cross-Adsorbed secondary antibody, Alexa Fluor™ 568 (Thermo Fisher Scientific, A10042) for 1 hour in the dark. The cells were then stained with 1μg/ml DAPI (Sigma, D9542) for 5 minutes in the dark. After washing, the coverslips were mounted in the glass slide using ProLong™ Gold Antifade Mountant (Invitrogen, P10144). The cells were visualized using the Carl Zeiss Axio Observer fluorescence microscope and the images were captured using ZEN pro software.

### Quantification of Macc1 expression

Cells were imaged at 63x magnification with respect to the nuclear plane with same imaging parameters across the group. The images were converted to grey scale. The blue (DAPI) and red (MACC1) channels were split separately. ROI was carefully drawn around well isolated cells. Integrated density (IntDen) of the cell was measured using ImageJ software for a minimum of 70 cells per group.

### RNA isolation and RT-qPCR

Total RNA was extracted using the Trizol® reagent (Thermo Fisher Scientific, 38229090), and 1.5 µg of RNA was used to synthesize cDNA using High-Capacity cDNA Reverse Transcription Kit (Thermo Fisher Scientific, 4374966). Real-time PCR was performed using iTaq Univer SYBR Green (BioRad,1725121) on the QuantStudio 7 Flex RealTime PCR System (Biorad, CFX96 Touch Real-Time PCR Detection System). The primers used are tabulated in Supplementary Table 4. The PCR conditions included an initial step at 95°C for 30 seconds for cDNA activation, followed by 39 cycles of 95°C for 5 seconds (denaturation) and 60°C for 30 seconds (annealing/extension). A melt curve analysis was performed from 65°C to 95°C, with 5-second increments per step. All reactions were carried out in triplicate, and transcript levels were normalized to β-actin. Relative fold change was calculated using the equation 2^−ΔΔCT^. Statistical analysis was conducted using two-way ANOVA in GraphPad Prism.

### Cell cycle analysis

Cells were seeded into a 24-well dish and were treated with Lovastatin (8.7μM) for 48 hours. The cells were harvested and washed twice with 1X DPBS, fixed with ice-cold 70% ethanol at -20 ° C for an hour and stained with propidium iodide staining solution in PBS 1X (0.1% Triton ™ X-100 (Sigma), 100 μg/mL PureLink™ RNase A (Invitrogen, 12091-021) and 50 μg/mL propidium iodide (Invitrogen, P1304MP) for an hour. The cell cycle distribution was determined using flow cytometer (BD FACS Verse, BD Biosciences) and the data was analyzed using FlowJo version 10 software. A minimum of 10,000 events were recorded for each cell line.

### Protein-Ligand Docking

To investigate the interaction between the transcription factor Sp1 and lovastatin, we first queried the RCSB Protein Data Bank (PDB) for any experimentally resolved full-length structures of Sp1. However, no such structures were available. Only 11 NMR structures corresponding to the C2H2 zinc finger domains (types 1, 2, and 3) were found. Given this limitation, we turned to the AlphaFold Protein Structure Database and retrieved the predicted structure of Sp1 (AF-P08047-F1-v4). Upon evaluation, we observed that a significant portion of the predicted structure had low model confidence scores (pLDDT < 50), rendering them unreliable for molecular docking. To ensure structural accuracy, we selected only the regions with high model confidence (pLDDT > 70). This filtered segment corresponded to residues 625–719, which encompass the type 1, 2, and 3 zinc finger motifs essential for DNA binding and potential ligand interaction.

a. Cavity Detection: The initial step in the docking workflow involved the identification of potential binding pockets on the selected Sp1 segment. For this purpose, CB-Dock2 was utilized, which implements a curvature-based method combining geometric and physicochemical analyses to detect concave surface regions that could accommodate small-molecule ligands. This cavity detection step was critical in guiding the subsequent docking simulations by narrowing down candidate binding sites based on their structural and chemical properties.
b. Template-Independent Docking: Following cavity detection, we performed template-independent docking using AutoDock Vina within the CB-Dock2 platform. This approach allowed unbiased modeling of the interactions between Sp1 and lovastatin, without relying on predefined binding templates. AutoDock Vina uses an empirical scoring function and a stochastic global search algorithm to predict optimal ligand conformations within the protein’s binding pocket. The potential binding modes were ranked based on the Vina score (expressed in kcal/mol), which reflects the estimated binding affinity-the more negative the score, the stronger the predicted interaction. The docking pose with the lowest Vina score was selected for further analysis, as it represents the most energetically favorable conformation of the Sp1-lovastatin complex (35).
c. Chimera for Visualization and Analysis: Chimera, a powerful molecular visualization software, was used for the visualization and analysis of protein structures and docked complexes (34).

### Combination therapy

Cells were seeded in triplicates in a 96 well plate. After 24 hours, lovastatin and 5-FU was added as monotherapy or in combination to Bulk, T1 and L1 cells with the concentration corresponding to the IC50 of the Bulk tumor. The cell viability was measured post 48 hours of treatment using MTT assay.

## Data Availability Statement

The data that support the findings of this study are available from the corresponding author upon request.

