## Supplementary Figures for "Repurposing statins in combination therapy for effective ablation of metastatic breast cancer"

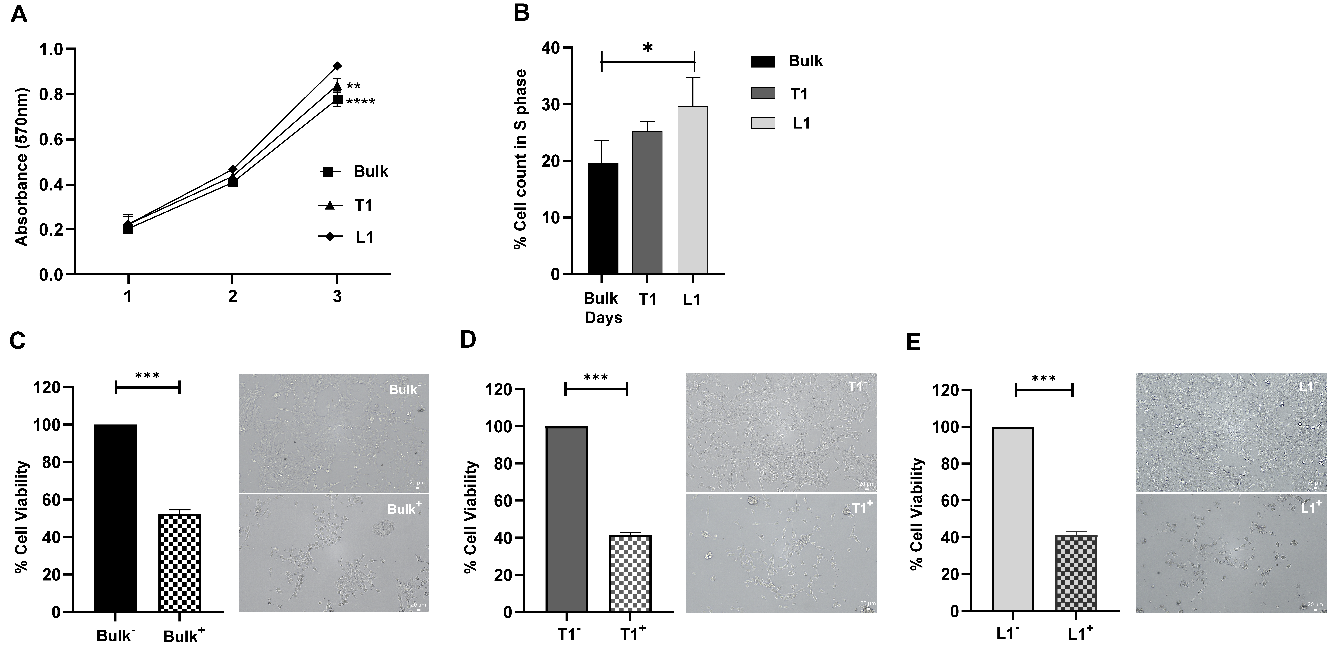

**Figure S1.** Phenotypic characterization and effect of lovastatin on metastatic tumor cells A) Proliferation analysis of Bulk, T1 and L1 cells using MTT assay indicated that L1 cells proliferate at a faster rate as compared to the Bulk and T1 cells. B) Cell cycle analysis of Bulk, T1 and L1 cells revealed a significant increase in the S phase of L1 cells as compared to the Bulk. All the experiments were performed in three biological replicates and the data were expressed as mean ± standard deviation. One-way ANOVA statistical test was used. *p < 0.05, **p < 0.01, ***p < 0.0001. Effect of Lovastatin on the cell viability of C) Bulk, D) T1 and E) L1 cells. Statistical significance was calculated using Unpaired t-test with Welch's correction with *** p value< 0.0002 and **** p value <0.0001.

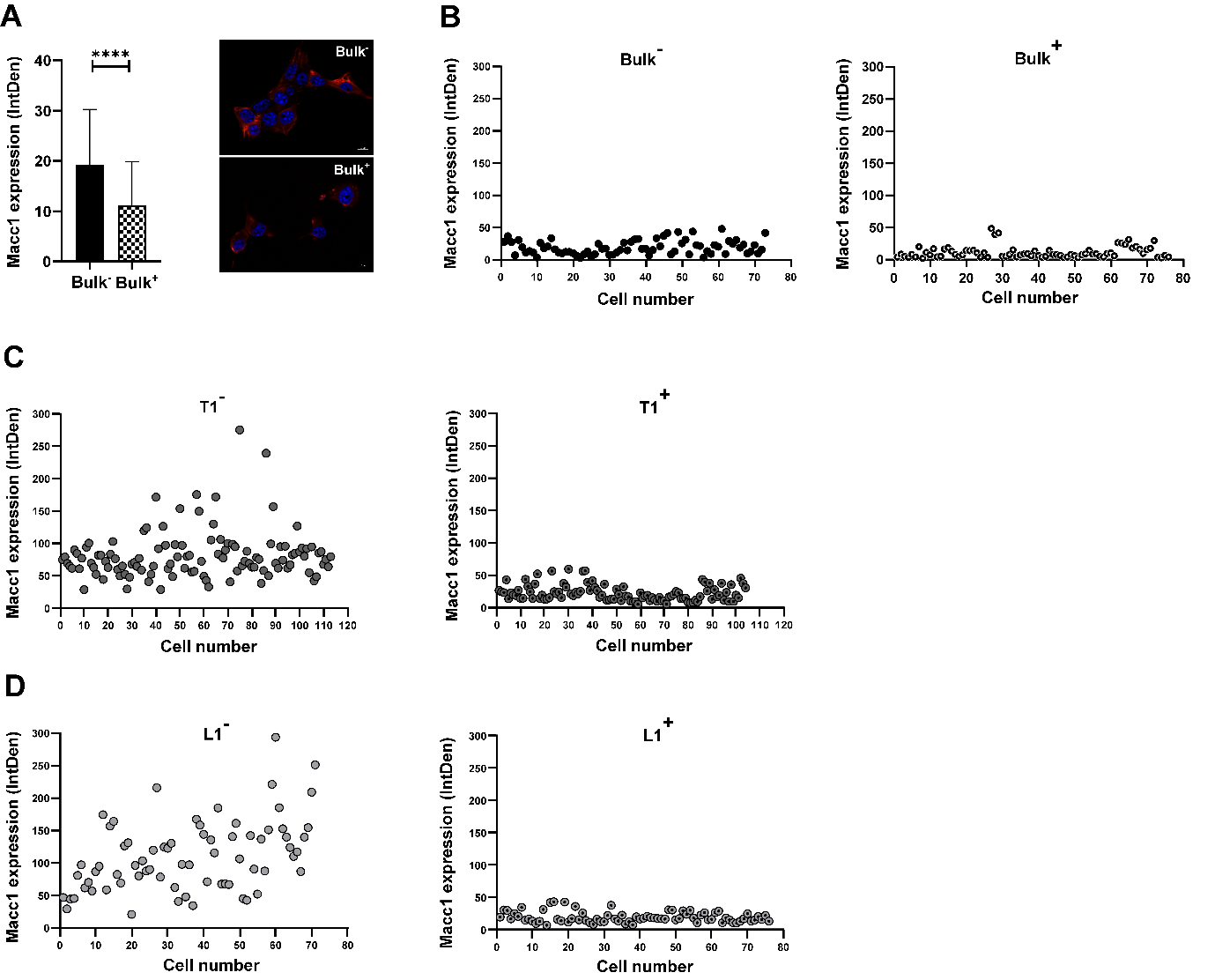

**Figure S2.** Effect of lovastatin on Macc1 expression of metastatic tumor cells. A) Effect of lovastatin on the Macc1 expression in the Bulk tumor cells. Representative immunofluorescence images showing the Macc1 expression in untreated and lovastatin treated groups of Bulk tumor cells. Scatter plots showing the Macc1 expression in the untreated and treated group at single cell level in B) Bulk, C) T1 and D) L1 cells. Statistical significance was calculated using Unpaired t-test with Welch's correction with *** p value< 0.0002 and **** p value <0.0001.

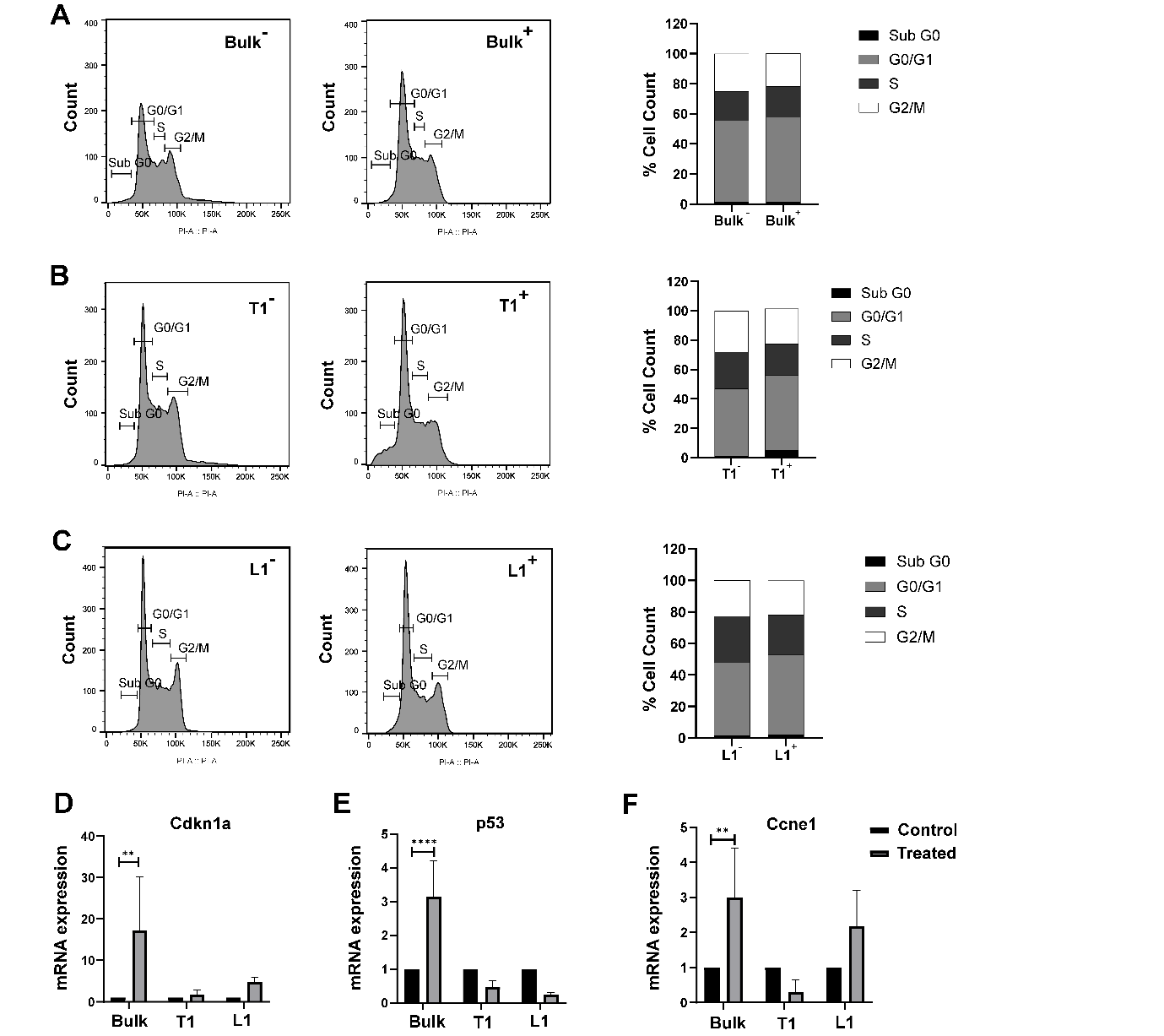

**Figure S3.** Effect of lovastatin on cell cycle phases in Bulk, T1 and L1 cells. Histogram and stacked bar graph depicting the cell cycle phases and the percentage cell count in in the A) Bulk B) T1 and C) L1 cells treated with lovastatin. The mRNA expression levels of the cell cycle genes D) Cdkn1a E) Tp53 and F) Ccne1 in untreated and treated Bulk, T1 and L1 cells using qRT-PCR. The data was normalized to β-actin and analyzed using 2^-ΔΔ^*^CT^* method. All the experiments were performed in three biological replicates and the data were expressed as mean ± standard deviation. Two-way ANOVA statistical test was used. *p < 0.05, **p < 0.01, ***p < 0.0001.

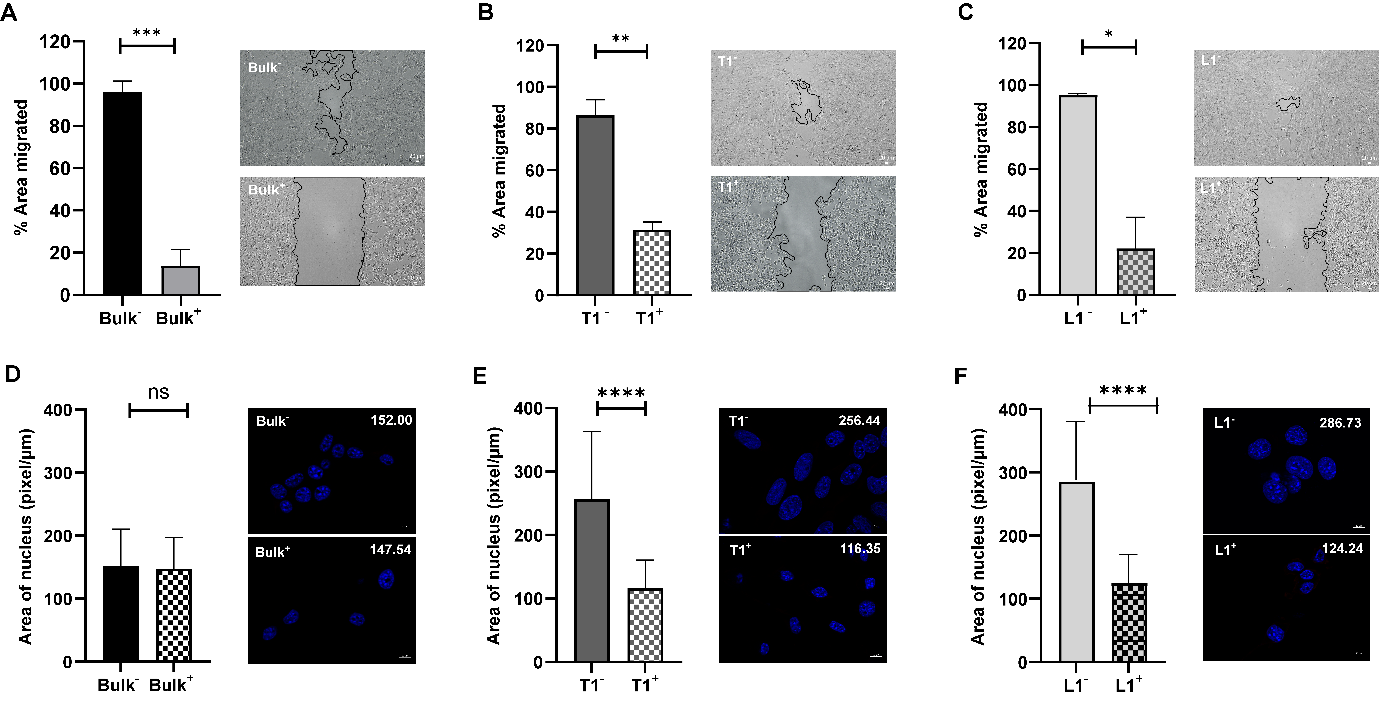

**Figure S4**. Pleiotropic effect of lovastatin on metastatic tumor cells. Lovastatin affects migration ability of the A) Bulk B) T1 and C) L1 cells upon 24 hours of Lovastatin treatment at 8.7 µM concentration. Quantification of the nuclear size of the D) Bulk E) T1 and F) L1 cells upon 48 hours of Lovastatin treatment at 8.7 µM. Representative images of cells stained with DAPI taken at 63x magnification with a corresponding scale bar of 10 µm. A minimum of 70 cells were quantified per group. Statistical significance was calculated using Unpaired t-test with Welch's correction. ns- non-significant, *** p < 0.0002 and **** p <0.0001.

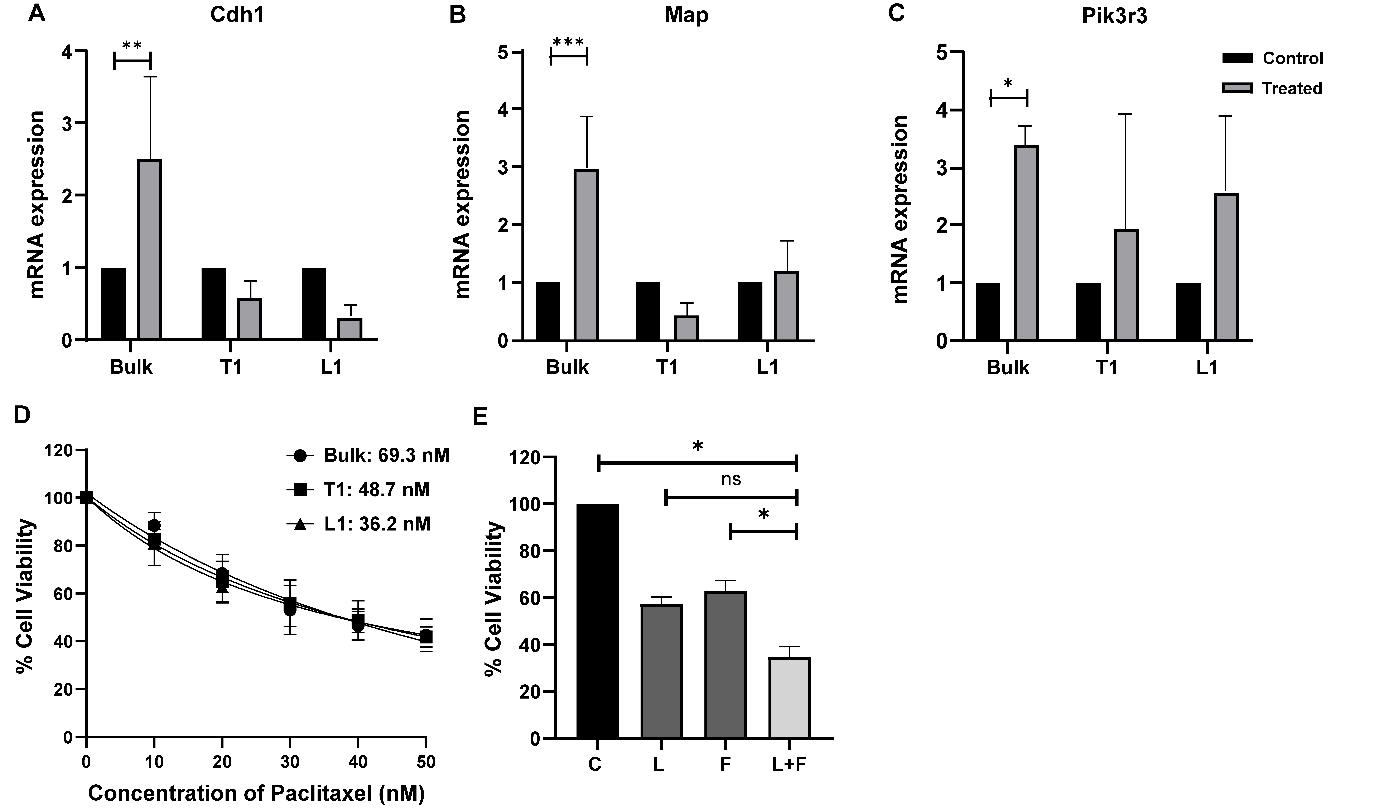

**Figure S5.** Key genes involved in molecular signaling pathways associated with Macc1 expression were analyzed using RT-qPCR. A) Cdh1 (E-cadherin), B) Map and C) Pik3r3 expression significantly increased in the Bulk tumor cells, but showed no significant change in T1 and L1 cells. D) Graph representing the dose-dependent effect of paclitaxel on Bulk, T1 and L1 cells post 48 hours of treatment. T1 and L1 cells did not display resistance to paclitaxel when compared to the Bulk. E) Graph representing the effect of 5-FU in combination with Lovastatin in the Bulk tumor cells. The data was normalized to β-actin and analyzed using 2^-ΔΔ^*^CT^* method. All the experiments were performed in three biological replicates and the data were expressed as mean ± standard deviation. Two-way ANOVA statistical test was used. *p < 0.05, **p < 0.01, ***p < 0.0001.
