## Supplementary Tables for "Repurposing statins in combination therapy for effective ablation of metastatic breast cancer"

| Macc1 expression /  % of cells | Bulk ^-^ | Bulk ^+^ | T1 ^-^ | T1 ^+^ | L1 ^-^ | L1 ^+^ |
| --- | --- | --- | --- | --- | --- | --- |
| 0-50 | 100 | 100 | 13.27 | 95.19 | 14.08 | 100 |
| 50-100 | 0 | 0 | 70.80 | 4.81 | 36.62 | 0 |
| 100-150 | 0 | 0 | 9.73 | 0.00 | 26.76 | 0 |
| 150-200 | 0 | 0 | 4.42 | 0.00 | 15.49 | 0 |
| 200-250 | 0 | 0 | 0.88 | 0.00 | 4.23 | 0 |
| 250-300 | 0 | 0 | 0.88 | 0.00 | 2.82 | 0 |

**Table S1.** Macc1 expression in Bulk, T1 and L1 cells, post-treatment with Lovastatin. Macc1 expression (IntDen) was binned into six categories ranging from 0 to 300. The percentage of cells in each category of Control and Lovastatin treated groups in Bulk, T1, and L1 cells is summarised in the table.

| Gene | Cells | Avg. Fold change | Regulation  (Treated vs Control) | Significance | p value |
| --- | --- | --- | --- | --- | --- |
| Cdkn1a | Bulk | 17.27 | ↑ | ** | 0.0027 |
|  | T1 | 1.75 | ↑ | ns | 0.8641 |
|  | L1 | 4.75 | ↑ | ns | 0.401 |
| Trp53 | Bulk | 3.13 | ↑ | **** | <0.0001 |
|  | T1 | 0.46 | ↓ | ns | 0.1653 |
|  | L1 | 0.24 | ↓ | ns | 0.0593 |
| Cdk4 | Bulk | 3.21 | ↑ | ns | 0.8452 |
|  | T1 | 0.9 | ↓ | ns | 0.9927 |
|  | L1 | 19.92 | ↑ | ns | 0.1143 |
| Ccne1 | Bulk | 3 | ↑ | ** | 0.0054 |
|  | T1 | 0.31 | ↓ | ns | 0.2622 |
|  | L1 | 2.18 | ↑ | ns | 0.0675 |
| Ccna2 | Bulk | 2.05 | ↑ | ns | 0.1951 |
|  | T1 | 1.09 | ↑ | ns | 0.9114 |
|  | L1 | 1.35 | ↑ | ns | 0.6515 |

**Table S2.** Cell cycle gene regulation in lovastatin-treated cells normalised to control. Relative mRNA expression levels of the cell cycle genes were measured by RT-qPCR. Cells were treated with lovostain and total RNA was extracted for quantitative PCR analysis. Gene expression levels were normalized to housekeeping gene β-actin and fold changes were calculated relative to the untreated control group (set as 1). Data represent mean ± standard deviation from independent experiments. Two-way ANOVA statistical test was used. *p < 0.05, **p < 0.01, ***p < 0.0001.

| Gene | Cells | Avg. Fold change | Regulation  (Treated vs Control) | 2-way Anova | |
| --- | --- | --- | --- | --- | --- |
|  |  |  |  | Significance | P value |
| Macc1 | Bulk | 0.2 | ↓ | **** | <0.0001 |
|  | T1 | 0.7 | ↓ | ** | 0.0021 |
|  | L1 | 0.06 | ↓ | **** | <0.0001 |
| Met | Bulk | 1.13 | - | ns | 0.7224 |
|  | T1 | 0.6 | ↓ | ns | 0.2693 |
|  | L1 | 0.96 | - | ns | 0.9994 |
| Wnt7a | Bulk | 1.6 | ↑ | ns | 0.0973 |
|  | T1 | 0.77 | ↓ | ns | 0.4953 |
|  | L1 | 1.11 | - | ns | 0.7534 |
| Pik3r3 | Bulk | 3.38 | ↑ | * | 0.0122 |
|  | T1 | 1.94 | ↑ | ns | 0.2684 |
|  | L1 | 2.57 | ↑ | ns | 0.076 |
| Map | Bulk | 2.98 | ↑ | *** | 0.0001 |
|  | T1 | 0.42 | ↓ | ns | 0.1246 |
|  | L1 | 1.21 | ↑ | ns | 0.5672 |
| Cdh1 | Bulk | 2.83 | ↑ | ** | 0.0026 |
|  | T1 | 0.72 | ↓ | ns | 0.304 |
|  | L1 | 0.49 | ↓ | ns | 0.1127 |
| Vim | Bulk | 2.51 | ↑ | * | 0.025 |
|  | T1 | 0.57 | ↓ | * | 0.048 |
|  | L1 | 0.32 | ↓ | ns | 0.0811 |
| Nr2f1 | Bulk | 1.4 | ↑ | **** | <0.0001 |
|  | T1 | 1.35 | ↑ | ns | 0.1439 |
|  | L1 | 0.7 | ↓ | * | 0.0151 |
| Tgfβ2 | Bulk | 0.56 | ↓ | * | 0.0375 |
|  | T1 | 0.48 | ↓ | * | 0.0165 |
|  | L1 | 0.36 | ↑ | ** | 0.0048 |

**Table S3**. Macc1 molecular pathway gene regulation in lovastatin treated cells normalised to control. Relative mRNA expression levels of the cell cycle genes were measured by RT-qPCR. Cells were treated with lovostain and total RNA was extracted for quantitative PCR analysis. Gene expression levels were normalized to housekeeping gene β-actin and fold changes were calculated relative to the untreated control group (set as 1). Data represent mean ± standard deviation from independent experiments. Two-way ANOVA statistical test was used. *p < 0.05, **p < 0.01, ***p < 0.0001.

| Gene | FP | RP |
| --- | --- | --- |
| Macc1 molecular pathway primers | | |
| β-actin (M) | AGAGGGAAATCGTGCGTGAC | CAATAGTGATGACCTGGCCGT |
| Macc1 (M) | ATTGACATGGAAGCCTCGAA | AGAAATGGGTTTGAGGCAGA |
| Met (M) | GCAGTGACGAGTATCGGACA | ATGGATGTCAGGAGCACTTGG |
| Wnt7a | GACAAATACAACGAGGCCGT | GGCTGTCTTATTGCAGGCTC |
| Pik3r3 | ACCACGAGTCTCTCGCTCAGTA | CCTGATACTGAGAGTGGAACTCC |
| Mapk13 | CAGCGAGGATAAGGTCCAGTAC | GCTCACAGTCTTCATTCACAGCC |
| Cdh1 (M) | GGTTTTCTACAGCATCACCG | GCTTCCCCATTTGATGACAC |
| Vim (M) | CCGTTGAAGGTCAAGACGTGCCA | AGGAGGCCGAAAGCACCCTGC |
| Nr2f2 (Dual) | ATTCTTCCTCGCTGAACCG | ACAGGAACTGTCCCATCGAC |
| TgfB2 (M) | GCAGATCCTGAGCAAGCTG | GTAGGGTCTGTAGAAAGTGGGC |
| Cell cycle primers | | |
| Cdkn1a/p21 (Dual) | GCAGATCCACAGCGATATCC | CAACTGCTCACTGTCCACGG |
| Trp53 (Dual) | AAGATCCGCGGGCGTAA | CATCCTTTAACTCTAAGGCCTCATTC |
| Cdk4 (Dual) | TATGAACCCGTGGCTGAAAT | TATGAACCCGTGGCTGAAAT |
| Ccne1 (Dual) | CTGAGAGATGAGCACTTTCTG | GAGCTTATAGACTTCGCACACCT |
| Ccna2 (Dual) | CAAGACTCGACGGGTTGCTC | GCTGGCCTCTTCTGAGTCTC |

**Table S4.** List of primer sequences used in RT-qPCR.
